# CellGPS: Whole-body tracking of single cells by positron emission tomography

**DOI:** 10.1101/745224

**Authors:** Kyung Oh Jung, Tae Jin Kim, Jung Ho Yu, Siyeon Rhee, Wei Zhao, Byunghang Ha, Kristy Red-Horse, Sanjiv Sam Gambhir, Guillem Pratx

## Abstract

*In vivo* molecular imaging tools are critically important for determining the role played by cell trafficking in biological processes and cellular therapies. However, existing tools measure average cell behavior and not the kinetics and migration routes of individual cells inside the body. Furthermore, efflux and non-specific accumulation of contrast agents are confounding factors, leading to inaccurate estimation of cell distribution *in vivo*. In view of these challenges, we report the development of a “cellular GPS” capable of tracking single cells inside living subjects with exquisite sensitivity. We use mesoporous silica nanoparticles (MSN) to concentrate ^68^Ga radioisotope into live cells and inject these cells into live mice. From the pattern of annihilation photons detected by positron emission tomography (PET), we infer, in real time, the position of individual cells with respect to anatomical landmarks derived from X-ray computed tomography (CT). To demonstrate this technique, a single human breast cancer cell was tracked in a mouse model of experimental metastasis. The cell arrested in the lungs 2-3 seconds after tail-vein injection. Its average velocity was estimated at around 50 mm/s, consistent with blood flow rate. Other cells were tracked after injection through other routes, but no motion was detected within 10 min of acquisition. Single-cell tracking could be applied to determine the kinetics of cell trafficking and arrest during the earliest phase of the metastatic cascade, the trafficking of immune cells during cancer immunotherapy, or the distribution of cells after transplantation in regenerative medicine.

## Introduction

*In vivo* molecular imaging tools are critically important to understand the role played by cell trafficking in physiological and pathological processes^1^. Metastasis, for instance, is the single most important factor for predicting cancer survival, and targeting the metastatic cascade could significantly improve clinical outcomes^2^. Other incurable diseases have seen unprecedented progress thanks to the development of cellular therapies^3–5^, using stem cells or immune cells to target specific disease processes. In addition, many physiological processes such as hematopoiesis, immunity and embryonic development rely on the precisely orchestrated migration of cells throughout the body^6,7^.

Molecular imaging tools can be used to estimate distribution of cells non-invasively, however, the information they provide relates to population averages and not individual cells. Unfortunately, many biological properties of cells become lost once averaged in this manner, including specific migration routes and kinetics of individual cells. Intravital microscopy methods can interrogate single-cell processes *in vivo*, but not at the scale of the whole body, and only in shallow tissues^8,9^. Additionally, magnetic resonance imaging (MRI) has demonstrated the capability to image single cells *in vivo*^10,11^, but only in uniform-background anatomical sites such as the brain and the liver, and without sufficient temporal resolution to track moving cells.

Positron emission tomography (PET) is the only non-invasive imaging modality capable of detecting picomolar probe concentration in humans. Owing to its exceptional sensitivity, PET has been applied to a variety of cell tracking applications^12–14^ but its potential for single-cell imaging has yet to be demonstrated. The concept of tracking moving particles using PET data originates from the field of chemical engineering, where the approach is used to estimate fluid and powder flows in complex opaque systems^15^. Recently, we developed a mathematical framework to extend this approach to the tracking of radiolabeled cells^16^, and validated the methodology experimentally using radioactive droplets^17^.

Here, we report the *in vivo* feasibility of “CellGPS”, a methodology for tracking the trafficking of single cells inside a living subject with exquisite sensitivity (Fig. 1). We use nanoparticles to stably and efficiently ferry ^68^Ga radioisotope into live cells, then isolate single radiolabeled cells for injection into live mice. From the pattern of high-energy photons detected by PET, we infer, in real time, the position of these single cells with respect to anatomical landmarks derived from X-ray computed tomography (CT).

**Figure 1.**
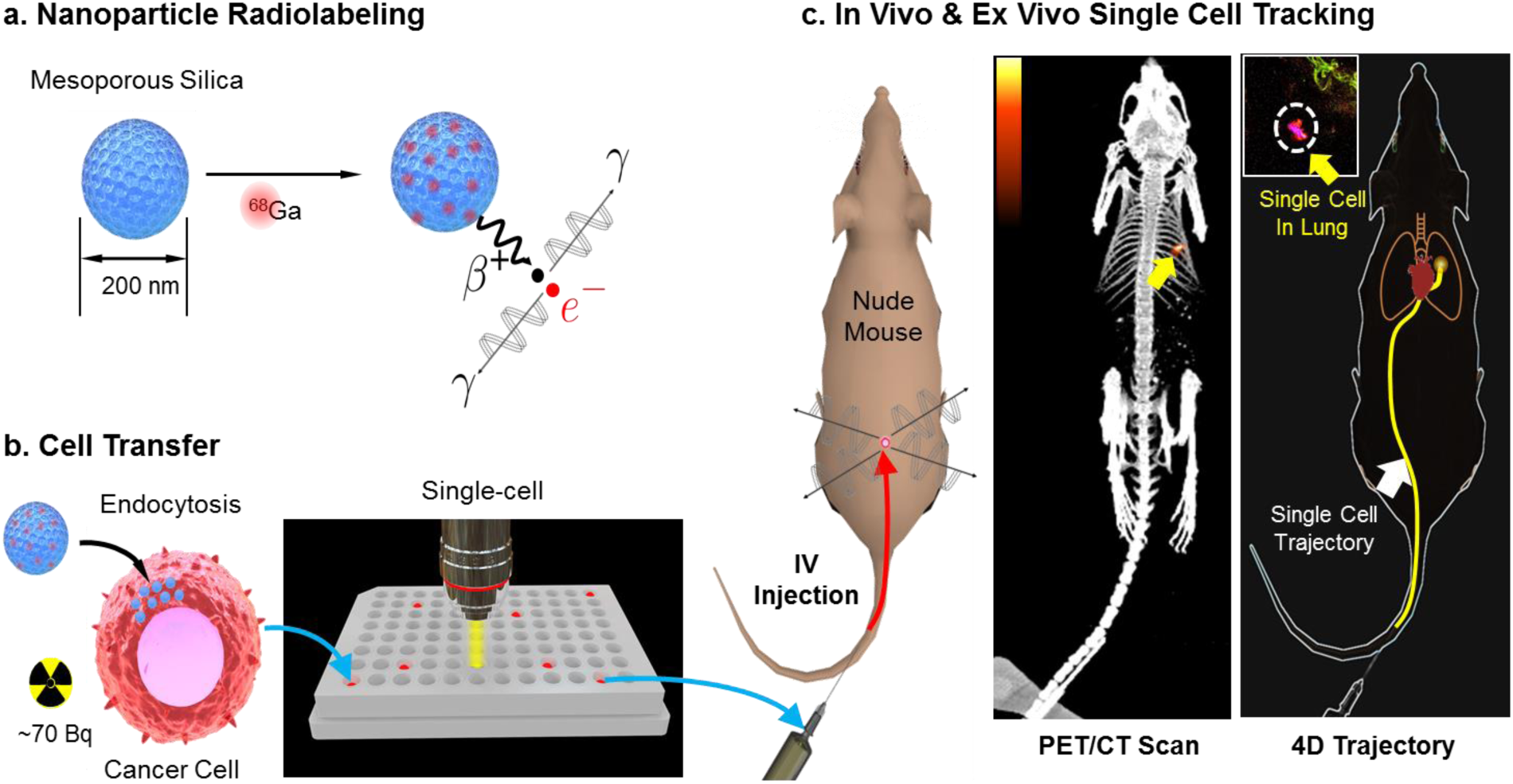
Overview of the CellGPS system. (a) Mesoporous silica nanoparticles (MSNs) are used to concentrate ^68^Ga, a clinical PET radioisotope with a 67 min half-life. (b) The nanoparticles are transferred to cells to achieve upwards of 100 Bq / cell (less than 1 millionth of a standard PET dose). (c) Isolated single cells are administered into mice. Gamma rays emitted from the single cell are captured by a small-animal PET scanner and processed by an algorithm to estimate the location of the moving cell in real time. In the example of human breast cancer cells administered IV, single cells arrested in the lung, as confirmed by *ex vivo* dissection.

## Results

### Efficient method for cell radiolabeling

To radiolabel live cells with high efficiency, we harness the exceptional cargo-carrying capacity of mesoporous silica nanoparticles (MSNs). Due to the large surface area of their pores, these nanoparticles can efficiently transfer radionuclides into cells for real-time cell tracking applications. This study specifically considers ^68^Ga, a radiometal used for PET imaging of neuroendocrine tumors and metastatic prostate cancer. Despite is relatively short half-life (67 minutes), high-specific-activity ^68^Ga can be eluted from inexpensive ^68^Ge generators, making it an increasingly popular isotope for clinical applications^18^. Here, we show that MSNs are able to carry ^68^Ga into live cancer cells to achieve upwards of 100 Bq per single cell, an amount of radioactivity sufficient for real-time tracking applications^16,17^. Although 100 Bq is truly a minute amount of radioactivity (it is less than 1 millionth of a standard PET dose), in terms of concentration, 100 Bq/cell is equivalent to 50 GBq/mL, about 100 times the concentration of ^68^Ga eluted from a medical generator. The use of MSNs is therefore required to concentrate ^68^Ga into cells and achieve sufficient labeling.

MSNs were sourced from Sigma-Aldrich (St Louis, MO) and physically characterized to reveal a spherical morphology with pore structure clearly visible on transmission electron micrographs (Fig. 2a). Hydrodynamic size distribution was estimated as 100-300 nm diameter (**Supplementary Fig. 1a**).

**Figure 2.**
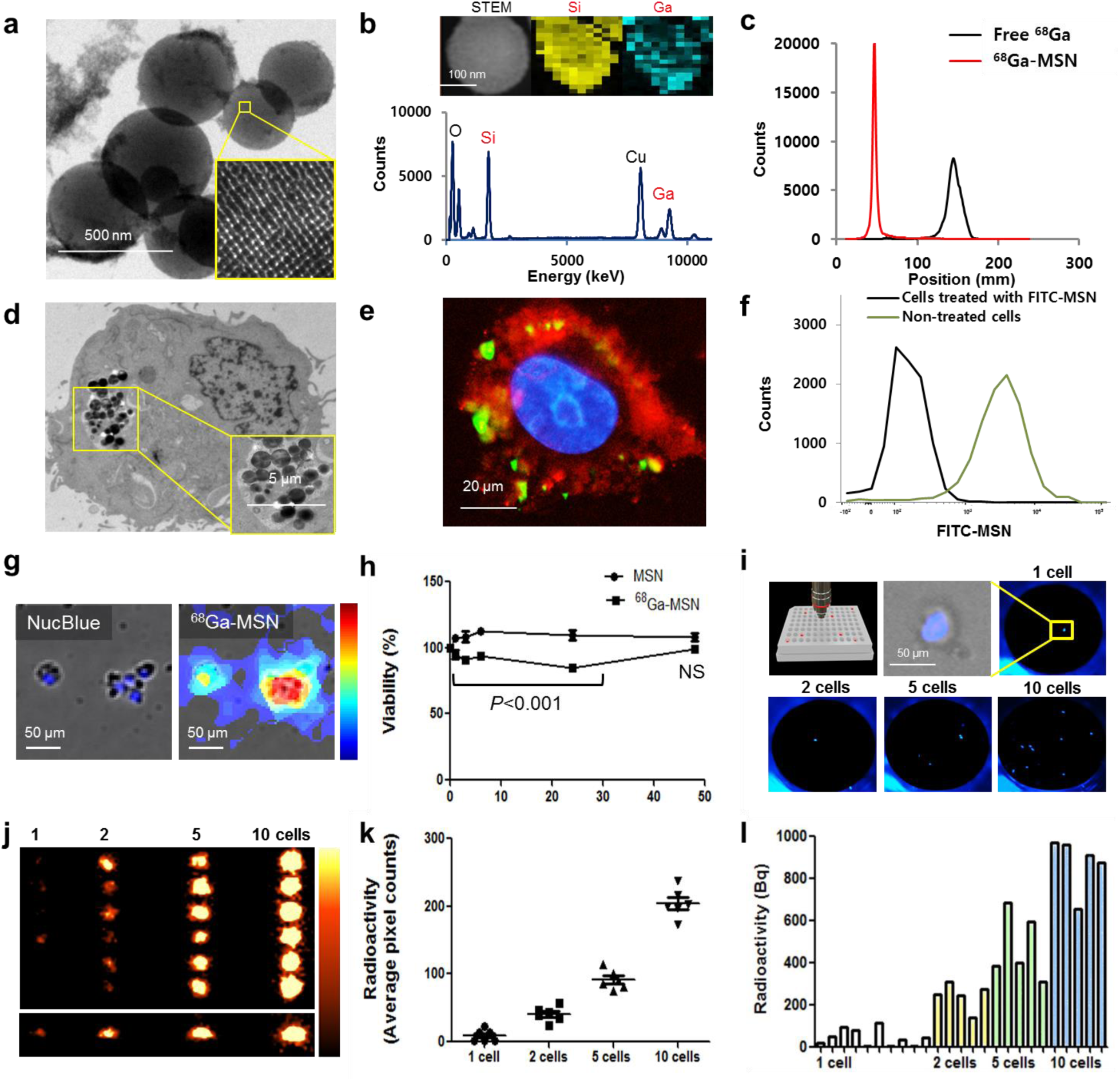
^68^Ga-MSN labeling method for sensitive cell tracking. (a) Transmission electron microscopy (TEM), showing MSN pore structure. (b) Energy dispersive spectroscopy and elemental mapping of single MSN labeled with stable ^69^Ga. (c) Rad**i**ochemical purity of ^68^Ga-MSN quantified using iTLC. (d) TEM showing uptake of lipid-coated MSNs by MDA-MB-231 cell. (e) Cellular uptake of FITC-labeled MSNs (green). Red is cell membrane and blue is nucleus. (f) Cell uptake of FITC-labeled MSN quantified by flow cytometry analysis. (g) Fluorescence (NucBlue) and radioluminescence microscopy (RLM) of MDA-MB-231 cells after uptake of ^68^Ga-MSN. (h) Cell viability assay following treatment with non-radioactive MSN and ^68^Ga-labeled MSN. Experiment performed in triplicate. NS, not significant. (i) Cell enumeration in Terasaki plates by fluorescence microscopy. (j) PET imaging of various numbers of ^68^Ga-MSN-labeled cells (top: coronal view, bottom: sagittal view) and (k) region-of-interest image quantification. (l) Cell uptake measured by *in vitro* gamma counting.

The MSNs were then labeled with ^68^Ga according to a newly reported chelator-free method that exploits the oxophilicity of most radiometals^19^. To optimize the method, MSNs were reacted with (non-radioactive) ^69^Ga under various conditions and the products were analyzed by electron microscopy (**Supplementary Fig. 1b**). Energy dispersive spectroscopy measurements were used to map the elemental composition of the labeled nanoparticles and found that ^69^Ga was uniformly distributed within the silica matrix (Fig. 2b). Other elements such as oxygen could be detected within the MSNs (**Supplementary Fig. 1c**).

The MSNs were then successfully labeled with clinical-grade ^68^Ga and purified to achieve >95% purity (thin-layer chromatography, Fig. 2c). Labeling efficiency, measured by gamma counting, was found to be as high as 0.5 Bq/nanoparticle. To enhance cell loading and biocompatibility, ^68^Ga-labeled MSNs were fused with a liposome-based transfection reagent^20^ to coat their surface with a lipid bilayer, as confirmed by electron microscopy (**Supplementary Fig. 1d**).

Passive uptake of ^68^Ga-labeled MSNs was demonstrated in MDA-MB-231 breast cancer cells. The cells were incubated with ^68^Ga-labeled MSNs (~25 MBq/ml) for 40 min, washed, then imaged by electron microscopy to reveal endocytosis of the particles into the cytosol (Fig 2d and **Supplementary Fig. 2a**). The MSNs were further labeled using a fluorescent dye (FITC), and their uptake was verified by fluorescence microscopy (Fig. 2e) and quantified by flow cytometry (Fig. 2f). Higher cell uptake was observed for lipid-coated MSNs compared to uncoated MSNs (**Supplementary Fig. 2b**). In quantitative terms, 95% of cells took up lipid-coated MSNs but only 20% took up uncoated MSNs (**Supplementary Fig. 2c** and **Supplementary Fig. 2d**).

Uptake of ^68^Ga-labeled MSNs by adherent MDA-MB-231 cells was confirmed by radioluminescence microscopy (RLM), a method with single-cell imaging resolution^21^. Areas of high radioactivity coincided with fluorescent nuclear staining (NucBlue) and brightfield cell imaging, indicating localization of radioactivity within cells (Fig. 2g). Cell labeling with ^68^Ga was more effective using lipid-coated MSNs than uncoated MSNs, and there was no detectable uptake of free ^68^Ga (**Supplementary Fig. 3a**). Furthermore, radiolabeling efficiency was relatively heterogeneous when analyzed on a single-cell basis. A few cells presented no detectable signal (**Supplementary Fig. 3b**). Radiolabel efflux by cells was measured by gamma counting and found to reach 50% after two hours (**Supplementary Fig. 4**). Finally, cell labeling using ^68^Ga-labeled MSNs resulted in a small (5-15%) but significant (*P*<0.001) decrease in cell viability for timepoints up to 24 h. At 48 h, cell damage was repaired, and no significant difference was observed (Fig. 2h). Non-radioactive MSNs did not significantly alter cell proliferation and viability.

Imaging weakly radioactive sources, such as single cells, is challenging because most PET scanners contain a small amount of naturally occurring radioactivity in their scintillation detectors^22^. Weak signals emitted by single cells may be drowned out by this intrinsic radioactivity background. We optimized the acquisition protocol and set the lower energy discriminator to 425 keV to achieve an optimal trade-off between background noise and sensitivity (**Supplementary Fig. 5**). With these settings, an Inveon PET/CT scanner achieved a background coincidence rate of ~ 2.6 counts/s, equivalent to ~40 Bq distributed throughout the field of view.

To demonstrate the feasibility of imaging single cells using PET, radiolabeled MDA-MB-231 cells were diluted and dispensed in Terasaki plates for enumeration (Fig. 2i). Wells containing 1, 2, 5, and 10 cells were identified by fluorescence microscopy and their contents were transferred to a standard 96-well plate for PET imaging. The images, which were reconstructed using the standard 2D-OSEM algorithm, showed increasing PET signal with increasing cell number (Fig. 2j). Importantly, a few single cells were clearly detectable by PET. Region-of-interest (ROI) analysis of the images confirmed the significant increase in ROI intensity with increasing cell number (Fig. 2k). Additionally, we performed quantitative gamma counting on another set of samples and measured a similar pattern of radioactivity uptake, with 30 Bq/cell on average and a range of 0-110 Bq/cell (Fig. 2l).

### Static PET imaging of small cell populations in mice

To determine whether single cells could be imaged *in vivo*, approximately 100 MDA-MB-231 cells were labeled with ^68^Ga-MSN (323 Bq in total) and DiO (fluorescent membrane dye), then injected via tail vein into a mouse. PET/CT images, acquired immediately thereafter, showed diffuse signal in both lungs (Fig. 3a) and little detectable signal in other organs (**Supplementary Fig. 6a**). *Ex vivo* gamma counting confirmed significant radioactivity in the lungs compared with liver and spleen (**Supplementary Fig. 6b**). To confirm the arrest of the cells in the lungs, lung tissues were excised, and DiO-labeled cancer cells were visualized by fluorescence microscopy (**Supplementary Fig. 6c & 6d**). The arrest of cancer cells in the lungs was confirmed in another mouse injected with 10^6^ DiO-labeled cells (**Supplementary Fig. 6e**).

**Figure 3.**
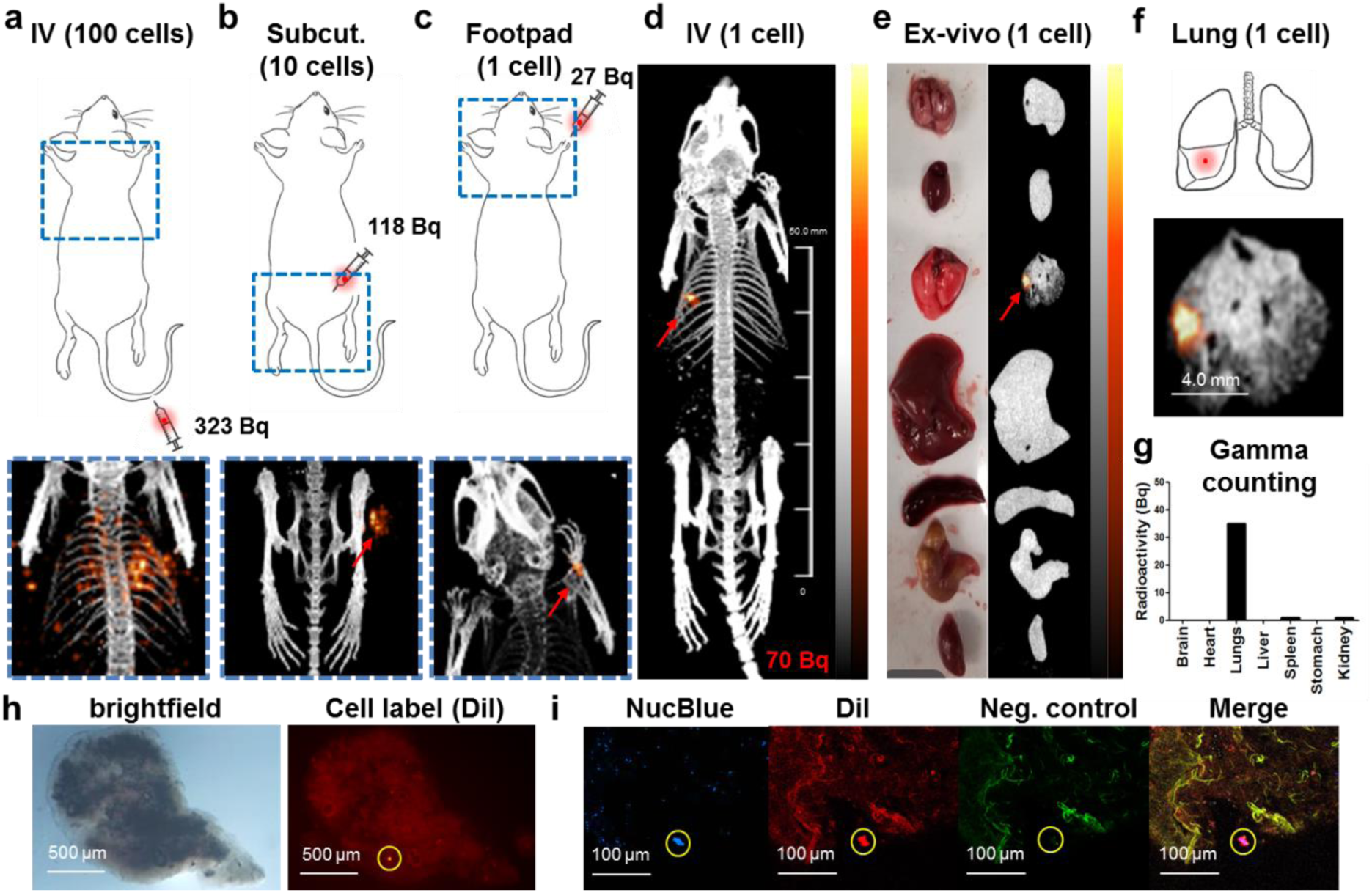
Static PET imaging of small cell populations in mice and *ex vivo* validation. (a) MDA-MB-231 cells arrested in the lungs following IV injection of 100 radiolabeled cells. (b) Visualization of a cluster of 10 cells after subcutaneous injection in the right thigh. (c) Single radiolabeled cell detected after its injection in the right forepaw. (d) Single radiolabeled cell after arrest in the left lung, following IV injection (*n*=3). (e) Photograph and PET/CT images of *ex vivo* organs from the same mouse (from top to bottom: brain, heart, lung, liver, spleen, stomach, kidney), and (f) magnified view of the lungs (*ex vivo*). (g) Gamma counting quantification of radioactivity of *ex vivo* organs. (h) Brightfield and fluorescence imaging of *ex vivo* lung tissue highlight single locus with high DiI fluorescence. (j) Confocal microscopy demonstrates dual staining for nucleus (blue) and membrane (DiI, red) but no green fluorescence (negative control), confirming the localization in the lung of the injected cancer cell.

Other injection routes were also investigated. A bolus of 10 radiolabeled cells (118 Bq in total) was injected in right thigh of a mouse and imaged with PET/CT images to reveal focal signal at the injection site (Fig. 3b), with no significant radioactivity elsewhere (**Supplementary Fig. 7a**). Similarly, 10 cells were injected intraperitoneally in the abdomen (94 Bq; **Supplementary Fig. 7b**) and in the footpad (84 Bq; **Supplementary Fig. 7c**). In all cases, the PET signal was confined to the injection site.

Using the same workflow, we proceeded to image single cells in mice using PET. A set of radiolabeled single cells was isolated in Terasaki plates through limiting dilution. Then, using a portable radiation survey meter, we rapidly screened dozens of cells to find single cells with high radioactivity that could be injected into mice. Because ^68^Ga-MSN uptake is variable, this screening process is critical to ensure that the candidate cell has sufficient radioactivity for *in vivo* tracking.

In a first experiment, a single radiolabeled single cell (27 Bq) was injected into the footpad of a mouse. PET/CT images, acquired minutes after injection, depicted strong focal signal near the injection site (Fig. 3c), with no significant PET signal in other organs (**Supplementary Fig. 7d**). These images show that PET can image single cells, which are much smaller than the resolution of the scanner. Indeed, the diameter of the PET signal was approximately 1.7 mm, which is consistent with the spatial resolution of the Inveon PET scanner.

Next, isolated single cells (20-70 Bq) were injected via a butterfly catheter into the tail vein of mice (*n*=3). PET acquisition was started either before (*n*=2) or after (*n*=1) single-cell injection. All injected mice presented a single locus of radiotracer accumulation in their lungs on PET/CT images (Fig. 3d). The major organs were then removed and imaged again *ex vivo* (Fig. 3e). Consistent with *in vivo* findings, focal uptake was observed in the lungs (Fig. 3f). *Ex vivo* gamma counting further measured 7-35 Bq in the lungs and less than 2 Bq in other organs (Fig. 3g). Finally, the lungs were dissected in an attempt to localize the arrested cancer cell. The tissue was chopped into five pieces; of these, only one contained significant radioactivity (**Supplementary Fig. 8**). We repeated this process until radioactive signal was confined to a piece of lung tissue smaller than 1 mm. We then imaged the tissue sample by widefield fluorescence microscopy and identified a single speck of enhanced red fluorescence (Fig. 3h). Finally, we imaged the area by confocal microscopy and found a single locus positive for blue and red fluorescence, which we attribute to NucBlue and DiI staining of the injected cell (Fig. 3i). Green fluorescence, imaged as a negative control, was negative for the identified locus.

### Dynamic tracking of single cells in mice

Cell trafficking is a dynamic process. Real-time tracking could help shed light on kinetics and migration routes of trafficking cells. Here, we demonstrate that PET data can be used to dynamically track the position of a single cell *in vivo*. By design, PET scanners record data in the list-mode format, a datafile that contains, in a long list, the attributes of every photon coincidence event detected by the scanner, including time of detection, photon energy, and position of the detector elements involved. In the conventional PET workflow, list-mode data are binned into a stack of 2D sinograms, then reconstructed to form a tomographic image using an algorithm called OSEM (ordered-subset expectation-maximization). In general, this approach does not permit dynamic tracking of single cells. Various particle tracking methods have been developed to estimate the time-varying position of a moving source directly the raw list-mode data, but their application has been limited to chemical engineering systems^15^. Our recent work modeled the trajectory of the cell as a cubic B-spline and computed its time-varying position by solving a convex optimization problem^16^.

The feasibility of tracking a moving cell was demonstrated first by enclosing a single radiolabeled MDA-MB-231 cell (36 Bq) into a 1.5 ml conical tube. Static PET/CT imaging detected a radioactive signal from the single cell inside the tube (**Supplementary Fig. 9a**). A list-mode acquisition was then performed, during which the tube was slowly moved along the axis of the scanner in one direction, then in the opposite direction, and finally removed from the scanner bore. The algorithm was able to compute the position of the object, even as it was being moved (**Supplementary Fig. 9b** and **Supplementary Video 1).** Furthermore, the count rate (**Supplementary Fig. 9c)** dropped (from 6.4 to 4.7 counts/s) once the vial had been removed from the scanner, suggesting that the cell contributed approximately 1.7 counts/s to the overall PET count rate, equivalent to 24 Bq (assuming 7% coincidence detection sensitivity).

To further validate this approach, we coiled a length of plastic tubing around a 3D-printed cylinder to form a helical test phantom (Fig. 4a). The phantom was placed in the bore of a PET scanner and a single MDA-MB-231 cell (67 Bq) was flowed at a speed of 1.36 mm/s through the tubing using a syringe pump. List-mode PET data were acquired for 10 min and then reconstructed into a 3D spline trajectory. Unlike the conventionally reconstructed image, which showed no localized signal (**Supplementary Fig. 10a**), the dynamically reconstructed trajectory matched the helical shape of the phantom, including its pitch and diameter (Fig. 4b). The single cell could also be seen exiting the scanner through the center of the phantom (**Supplementary Video. 2**). We quantified the localization error (defined here as the root-mean square deviation from the ideal helical trajectory) and found it to be less than 5 mm (**Supplementary Fig. 10b**). Finally, we estimated the velocity of the cell according to its reconstructed trajectory (Fig. 4c). At its peak, the cell moved at the same speed as the injected bolus, although, at times, the cell slowed down significantly, possibly because it interacted with the walls of the plastic tubing.

**Figure 4.**
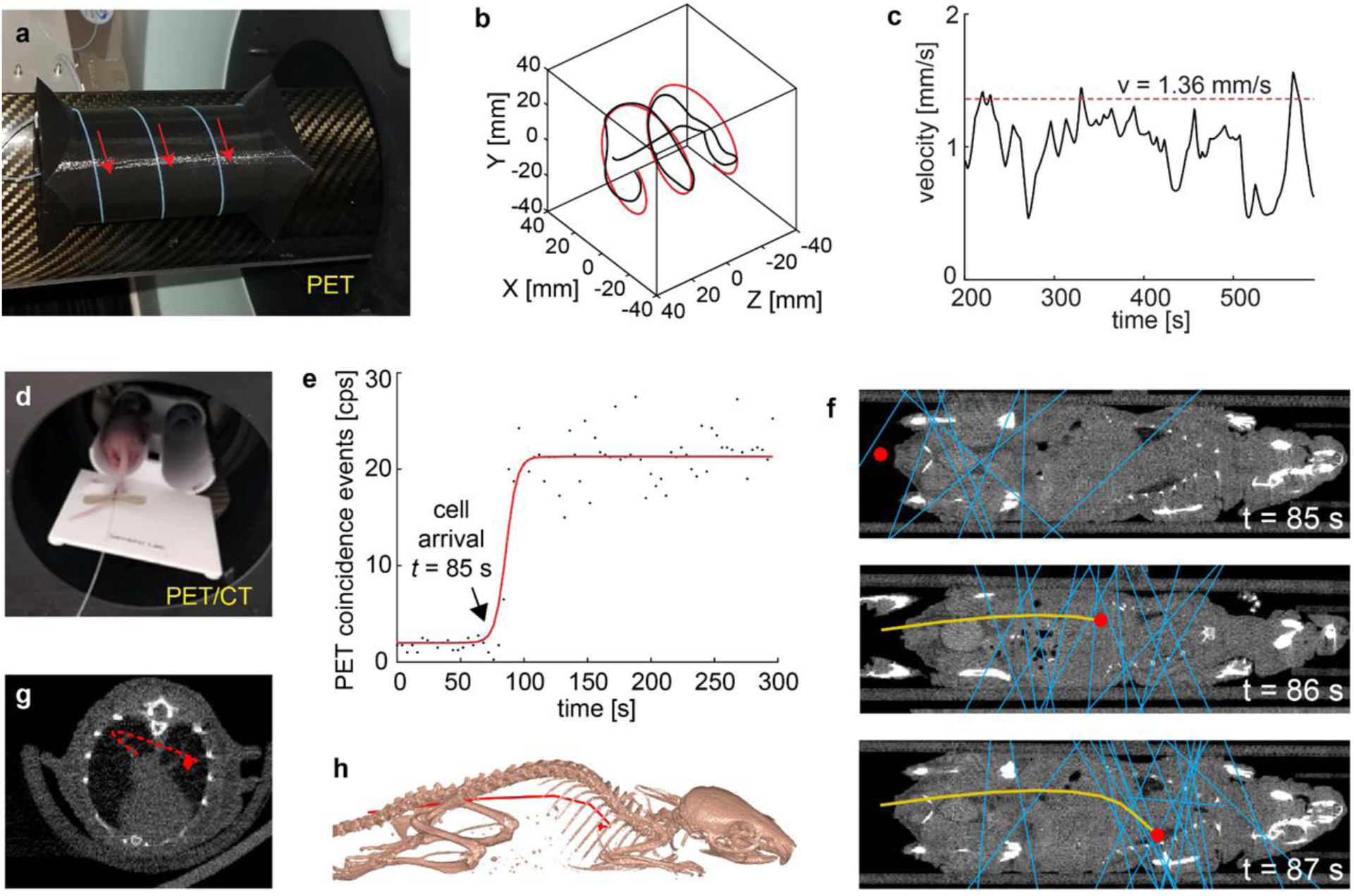
Dynamic PET tracking of single cells. (a) A single radiolabeled cell is flowed through a length of tubing coiled around a 3D-printed cylinder. (b) Cell trajectory reconstructed from list-mode PET data (black line) and closest helical fit (red line). (c) Estimated cell velocity as a function of time (black line), and average velocity of the cell medium (dashed red line). (d) Butterfly catheter installed into the tail vein for simultaneous cell injection and PET imaging. (e) Coincidence event rate recorded by the scanner. Events measured before cell arrival are due to background noise. (f) Dynamic tracking of single cell *in vivo*, shown at three different time points. The blue lines represent coincidence events detected by the scanner. The red circle is the estimated location of the cell. The yellow line is the reconstructed trajectory. Coronal CT views are shown for anatomical reference. (g) Reconstructed trajectory shown as a transaxial view, including cross-sectional CT slice through cell arrest location. (h) Reconstructed trajectory shown relative to 3D surface-rendered bony anatomy.

We then applied this dynamic tracking methodology to single cells injected into live mice. First, the trajectories of static cells were reconstructed. For the previous mouse injected in the footpad, the single cell (27 Bq) remained at the injection site for the duration of the 10 min PET acquisition (**Supplementary Fig. 11** and **Supplementary Video. 3**). For another mouse injected intravenously before the start of PET/CT acquisition, the radiolabeled cell (30 Bq) was correctly located in the mouse lung and no significant motion was observed during the 10 min acquisition (**Supplementary Fig. 12** and **Supplementary Video. 4**), suggesting that the cell did not migrate after its arrest in the lung. For these static cells, the 3D position reconstructed using the spline-based tracking algorithm matched the PET images reconstructed through conventional OSEM.

Finally, real-time tracking was demonstrated by tracking a single cell (70 Bq) continuously, from its injection into the tail vein (via a catheter; Fig. 4d) to its arrest in the lungs. We monitored the arrival of the radiolabeled cell into the field of view of the scanner by tracking the rate of coincidence events as a function of time (Fig. 4e). Initially, the detected counts were due to natural background radioactivity. Once the cell entered the bore of the scanner (at *t* = 85 s), the count rate jumped from 2 cps to 21 cps, and cell tracking was initiated. To visualize the pattern of coincident gamma ray detection, we charted the recorded coincidence events (shown as blue lines) and the reconstructed cell trajectory (yellow curved line) over a cross-sectional CT image of the mouse (Fig. 4f). The dynamic migration of the cell is also shown in **Supplementary Video 5** and **6**. According to the reconstructed trajectory, the single cell arrested in the lungs within 3 s of intravenous injection. The location of the arrested cell matched the focal signal seen on conventional PET/CT imaging (**Supplementary Fig. 13a**). The reconstructed cell migration route is also displayed as a transaxial view (Fig. 4g) and combined with a surface-rendered model of the mouse bony anatomy (Fig. 4h). An analysis of the reconstructed trajectory estimated the velocity of the cell at around 50 mm/s immediately post-injection (**Supplementary Fig. 13c**). For reference, the velocity of venous blood in the inferior vena cava in mice is approximately 40 mm/s, according to a previous study using magnetic resonance imaging^23^. Finally, as a control, PET data acquired from *ex vivo* organs were reconstructed. The algorithm localized the single cell to the excised lungs, and no motion was observed, apart from random fluctuations due to physical noise (root-mean-square error, RMSE <0.45 mm; **Supplementary Fig. 13d** and **Supplementary Video 7**).

## Discussion

One of the major findings of this study is that PET is sufficiently sensitive to image static single cells labeled with 20 Bq/cell or more of radioactivity. Standard OSEM reconstruction produced surprisingly good images of single cells *in vivo* after 10 min of acquisition with an optimized energy window that removes background noise. Labeled cells manifest in PET images as small hot spots (diameter ~1.7 mm) in an otherwise blank background. One advantage of tracking individual cells (as opposed to bulk populations) is that the presence of cells in an organ can be assessed unambiguously because, unlike single cells, unbound tracer does not produce focal uptake. Different numbers of cells could be imaged with this approach, but single-cell enumeration will only be possible where cells are sufficiently distant from one another. More advanced image reconstruction methods may provide advantages for single-cell imaging and should be investigated.

The second important finding of this study is that single migrating cells can be tracked dynamically directly from list-mode PET data. From a technical standpoint, dynamic cell tracking is vastly more challenging than static cell imaging, especially at high velocities. Our approach infers the position of the cell by fitting a 3D cubic B-spline to the dynamic pattern of gamma ray interactions recorded by the PET scanner. The method is flexible and can be tuned to achieve either high accuracy or high reactivity, which are two competing objectives. For instance, in Fig. 4 d-h, a single cell (70 Bq) was tracked from the tail vein to its arrest 3 s later in the lungs. During the first 5 s of tracking, the algorithm was tuned to be highly reactive to cell motion to accommodate the high velocity of the cell. Previous analyses indicate that robust tracking is achievable on the condition that ratio of the cell velocity (*V*) to its activity (*A*) does not exceed 0.1 mm/s/Bq^17^. Physically, *V*/*A* represents the average distance traveled by the cell in between radioactive decays. In this example, the initial velocity of the cell was estimated to be around 50 mm/s. Based on the *V*/*A* criterion, a single-cell activity of 500 Bq—considerably higher than 70 Bq–would be required for accurate tracking. The reconstructed trajectory of the cell during the first 5 s of tracking should therefore be interpreted as qualitative rather than quantitative. Once the cell arrested in the lungs, the algorithm was adjusted to maximize localization accuracy. With these settings, the variability in the estimated position of the cell over time was ~ 0.4 mm RMSE, considerably better than the spatial resolution of the scanner. The phantom data (Fig. 4 a-c) also illustrate the importance of the *V*/*A* criterion. In that experiment, a *V*/*A* ratio of 0.02 mm/s/Bq was achieved, enabling robust dynamic tracking of the moving single cell.

For greater cell throughput, the CellGPS methodology could be extended to enable tracking of multiple cells simultaneously. A previous study simultaneously tracked, *in vitro*, 16 yeast cells labeled with ^18^F (32 Bq/cell). Individual cells could be resolved and slow diffusion of the cells in the medium was observed^24^. Tracking multiple cells will likely be possible provided that the following conditions are met: cells are separated from each other by at least 2-3 times the spatial resolution of the scanner; at any given time, no more than one cell moves rapidly within the imaged subject; and cells are administered one-by-one such that they can be individually tracked from the injection site to the initial site of arrest. In the case of cellular therapies, where millions of cells are injected, it may be possible to label and track a small subset of cells using the CellGPS methodology.

Robust tracking of single cells using PET requires scanners with low background and high sensitivity. Preclinical PET scanners have much higher sensitivity than human-sized scanners because their bore almost completely surrounds the subject. However, almost all PET scanners suffer from non-negligible background noise arising from natural radioactivity within their lutetium oxyorthosilicate (LSO) detectors. Improved tracking performance can be achieved using PET detectors built from scintillator materials that do not contain significant radioactivity, such, for instance, bismuth germanate (BGO). In a previous study, we assessed that the amount of radioactivity required per cell for tracking could be reduced three-fold through the use of low-noise BGO detectors^17^.

With a half-life of 67 minutes, ^68^Ga is suitable for short-term tracking applications. For long-term tracking, MSNs can be labeled with other radiometals, such as ^64^Cu (12 h half-life) and ^89^Zr (78 h half-life)^19^. However, these long-lived isotopes may cause higher radiotoxicity to the labeled cells and they may also affect the biological fate of the labeled cells. Most of the biological damage is wreaked by positrons, which upon emission move at relativistic speeds through cells and ionize biomolecules along their path. Fortunately, the physical range of positrons is much greater than the typical cell diameter, therefore, most of the damage is outside the labeled cells. To mitigate radiotoxicity, biological radioprotectants could be loaded into cells prior to radiolabeling. These compounds reduce oxidative damage by scavenging free radicals and reactive oxygen species.

In conclusion, CellGPS represents an entirely novel workflow for tracking individual cells *in vivo*. The methodology was used to show that MDA-MB-231 cancer cells injected intravenously in mice arrest in the lung within 3 s of their injection and travel at a velocity of up to 50 mm/s.

This novel workflow could have a number of significant applications. For instance, it could be used to investigate the early stages of the metastatic cascade. Considerable evidence suggests that tumor cells combine with platelets to form microemboli complexes in the vasculature and that these complexes play a critical role in the formation of metastases^25^. Another significant prognostic factor is the presence of clusters of circulating tumor cell (CTC) in the bloodstream^26^. Given that CTC clusters migrate 10 times more slowly than single CTCs through peritumoral vessels^27^, the CellGPS workflow could be used to determine patterns of cell arrest at the scale of whole subjects for different types of metastatic dissemination. Cell velocity within blood vessels, measured in vivo using PET, could become a novel biomarker linking cell arrest in specific organs to the expression of cell adhesion molecules in the vascular bed.

Beyond the study of cancer metastasis, cellGPS could be used to measure routes and kinetics of immune cell mobilization in response to injury. Finally, the method could help support clinical trials of cellular therapies for cancer immunotherapy and regenerative medicine. In recent years, PET scanners with extended field of view have been developed for total-body human PET imaging^28^. These highly sensitive systems could one day be used for translational applications involving tracking of single cells in humans.

## Acknowledgements

This work was funded in part by NIH grants 5R21HL127900 and T32CA1186810. Experiments were performed at the small-animal imaging facility, the Stanford radiochemistry facility, and the Stanford nano shared facility. The authors wish to gratefully acknowledge support and assistance from the following individuals: Dr. Mirwais Wardak, Carmen Azevedo, Travis Shaffer, Hongquan Li, Frezghi Habte, Arut Natarajan, Seung-Min Park, Tim Doyle, and Suyeon Lee. The ShowVol package by Dirk-Jan Kroon was used for surface rendering.

## Author contribution statement

KOJ performed *in vitro* and *in vivo* experiments. TJK performed physical tests and processed data. JHY characterized nanoparticles and optimized their radiolabeling. SR imaged *ex vivo* specimens. WZ reconstructed images. BH contributed to the design of the methods. GP reconstructed dynamic cell trajectory data. KOJ and GP designed the study and wrote the manuscript. KRH, SSG and GP supervised the study.

## Online Methods

### Cell Line and Cell Culture

The MDA-MB-231 cell line (human invasive ductal carcinoma) was obtained from Cell Biolabs in February 2016 and cultured in Dulbecco’s Modified Eagle Medium (DMEM) medium supplemented with 10% fetal bovine serum (FBS) and 1% antibiotic–antimycotic mix. The cells were cultured at 37°C and 5% CO2 in a humidified environment, up to a passage number of 20.

### Physical Characterization of MSNs

Propylamine-functionalized mesoporous silica nanoparticles (Sigma-Aldrich, St Louis, MO, USA; cat# 749265-1G) with 200 nm particle diameter and 4 nm pore size were physically characterized. The particles were deposited on 300-mesh carbon/formvar coated copper grids for imaging with a transmission electron microscope (JEM-1400 series 120kV, JEOL USA Inc., Pleasanton, CA, USA) equipped with a LaB6 emitter and a Gatan Orius 10.7-megapixel CCD camera. The elemental compositions of the MSNs after labeling with non-radioactive ^69^Ga was validated by energy-dispersive X-ray spectroscopy (EDS) and element mapping labeling using a transmission electron spectroscopy instrument (FEI Tecnai G2 3F20X-Twin microscope at 200 keV) equipped with a scanning TEM (STEM) unit with bright field/dark field detector, x-ray detector (EDS), and CCD camera. Finally, the concentration and hydrodynamic size distribution of the MSNs were determined by nanoparticle tracking analysis (NanoSight Ltd, Amesbury, UK).

### MSN Radiolabeling

The MSNs were labeled with ^68^Ga using a chelator-free reaction method^19^. First, the nanoparticles were activated in ethanol overnight. Then, 4.5 µg of MSNs were added to a 1.5 ml tube containing 100 µl of MES buffer (0.1 M, pH 7.3). Subsequently, MSNs were labeled with ^68^Ga by adding 1.5 ml of ^68^GaCl_3_ solution (~600 MBq) eluted with HCl (0.1 N) from a clinical generator. To maintain a reaction pH of 7.3, 30 µl of amonium hydroxide solution (Sigma-Aldrich, St Louis, MO, USA) was added, and the mixture was incubated at 70°C for 20 min. The nanoparticles were then washed three times with PBS and centrifuged at 14,000 rpm for 1 min to remove residual ^68^Ga. To determine radiolabeling purity (% fraction of radioactivity bound to nanoparticles), the radiolabeled MSNs (1.5 μl) were deposited on a silica-gel impregnated iTLC-SG paper and 50 mM EDTA (pH 5) was used as the elution solvent. The samples were then analyzed using a Bioscan AR-2000 radio-TLC plate reader (BioScan Inc., Washington, DC, USA).

### Lipid Coating of MSN

To enhance MSN uptake in cells, a cationic liposome transfection agent (Lipofectamine 2000, Invitrogen, California, USA) was also used to coat the surface of MSNs with a lipid bilayer. Two separate samples were prepared: one vial containing a mixture of 4.5 µg of MSN and 500 µl of Opti-MEM (Invitrogen, California, USA), and the other containing a mixture of 20 µl of Lipofectamine diluted in 500 µl of Opti-MEM. After 5 min of incubation at room temperature, the two samples were mixed together and incubated for additional 15 min at room temperature. The coated MSNs were then recovered by centrifugation.

### Fluorescent Labeling of MSNs and Cells

The MSNs were labeled with FITC (fluorescein isothiocyanate isomer I; Sigma-Aldrich, St Louis, MO, USA), via formation of thiourea bond with NH2 functional groups on the MSN surface. Approximately 1 mg of MSNs were mixed with a solution of 1 ml FITC (0.1 mg/ml) and incubated at room temperature for 1 hr. The mixture was then washed three times with fresh PBS and centrifuged at 14,000 rpm for 1 min to remove excess FITC. For validation purpose, cells were labeled by incubation at 37°C for 15 min with fluorescent lipophilic tracers such as DiO (Invitrogen, Carlsbad, CA, USA; green fluorescence; Ex: 484 nm, Em: 501 nm) or DiI (Invitrogen, Carlsbad, CA, USA; red fluorescence; Ex: 565 nm, Em: 594 nm). Additionally, nuclear staining was performed by incubation with NucBlue Live (ReadyProbes; Thermo Fisher Scientific, Waltham, MA, USA) for 5 min at 37°C.

### Fluorescent Imaging and Flow Cytometry of MSN Uptake in Cells

MDA-MB-231 cells (2× 10^5^ cells/plate) were incubated with MSNs or lipid-coated MSNs (4.5 μg/mL) for 40 min. To characterize cell uptake of MSNs, the treated cells were fixed with 2% glutaraldehyde. The cells were subsequently deposited on 300-mesh carbon/formvar-coated copper grids and imaged by TEM as previously described. Similarly, the uptake of FITC-labeled MSNs was imaged by fluorescence microscopy (EVOS FL, ThermoFisher Scientific, Santa Clara, CA, USA) and further quantified by flow cytometry (BD FACSAria Fusion sorter, BD Bioscience, San Jose, CA, USA).

### Cell Viability Assay

MDA-MB-231 cells were seeded in 96-well plates (2.5 × 10^3^ cells/well). To evaluate the cytotoxicity of MSNs in cells, cell viability was determined at various time points, up to 48 h after treatment with 4.5 μg/mL of lipid-coated MSNs (with or without ^68^Ga labeling). The cells were incubated with CCK-8 (Cell Counting Kit-8) solution (Sigma-Aldrich, St Louis, MO, USA) and, after 1 h, mean optical density (OD) was measured at 450 nm using a GloMax multi-detection system (Promega, Madison, WI, USA). Blank measurements were acquired and subtracted.

### Radioluminescence Microscopy

A custom-built radioluminescence microscope (RLM) was used to image the uptake of ^68^Ga-labeled MSNs by cells. The construction and use of the microscope is described in a previous publication^21^. Prior to the experiment, MDA-MB-231 cells (2× 10^5^ cells) were seeded on a tissue-culture-treated 35 mm glass-bottom dish (ibidi Labware GmbH, Germany) and incubated in a CO_2_ incubator for 24 h. The cells were then treated with ^68^Ga-labeled MSN or lipid-coated ^68^Ga-labeled MSN (~25 MBq/ml, suspended in 3 ml DMEM) for 40 min at 37°C. The cells were rinsed three times with PBS, then incubated with a nuclear fluorescent stain (NucBlue Live) for 5 min at 37°C. To perform RLM imaging, a CdWO_4_ scintillator crystal (MTI Corp., Richmond, CA, USA) was placed on top of the radiolabeled cells. Fluorescence images were first captured using the DAPI channel (excitation/emission: 358 nm⁄461 nm) to image the nuclei of adherent cells. RLM images were then acquired by setting the camera parameters to 1200 EM gain, 4×4 pixel binning, 150 ms exposure time, and 24,000 camera frames. Radioluminescence images were reconstructed using ORBIT (Optical Reconstruction of the Beta-ionization Track), a MATLAB toolbox developed in-house^29^. Radioactivity of each cell is expressed as counts per minute (cpm) and is calculated by enumerating the number of radioactive decays per cell throughout the hour-long acquisition. Radioactivity per cell was determined by region-of-interest analysis in MATLAB.

### Radioactivity Measurement using D-PET and Gamma Counter

Cells labeled with ^68^Ga-MSN were manually sorted into single cells through limiting dilution in Terasaki plates^30^. Fluorescence microscopy was used to enumerate the cells in each well. Wells containing 1, 2, 5, and 10 cells were identified and their contents transferred to a 96-well plate for PET imaging (Inveon D-PET, Siemens Preclinical Solutions, Knoxville, TN; 425-650 keV energy window, 10 min acquisition time). The radioactivity of single wells was assessed by region of interest (ROI) analysis using the Inveon Research Workplace (IRW) software. Another set of cells was prepared and measured by gamma counting (AMG, Hidex, Turku, Finland) with a counting time of 30 s per sample.

### Radiolabel Efflux

MDA-MB-231 cells (2× 10^5^ cells) were seeded on tissue-culture-treated 35-mm glass-bottom dishes and incubated in a CO_2_ incubator for 24 h. The cells were then treated with lipid-coated ^68^Ga-labeled MSNs (~25 MBq/ml, 3 ml in DMEM) for 40 min at 37°C, washed three times with PBS, incubated for another 30, 60, 90, or 120 min at 37°C, then washed again three times with PBS. Once all the cells had been washed, the radioactivity remaining in the samples was measured using gamma counting.

### Dynamic Cell Tracking in Helical Phantom using PET/CT

A cylindrical phantom with helical grooves was 3D printed (Z18; MakerBot Industries, Brooklyn, New York, USA), and PEEK tubing was wrapped along the grooves (51 mm diameter, 27 mm helical pitch). The phantom was placed inside the bore of a PET scanner (Inveon D-PET, Siemens pre-clinical solutions, Knoxville, TN, US). Using a syringe pump, a single radiolabeled single cell (67 Bq) was flown through the phantom at velocity of 1.36 mm/s while PET data were acquired (425-650 keV energy window, 10 min acquisition time). The list-mode data, which contained 2,165 coincidence events, was imported into MATLAB and the cell trajectory was reconstructed using our previously reported algorithm^16^. This reconstruction algorithm models the trajectory of the cell as a 3D cubic B-spline. The number of knots was set to have approximately 30 events per spline knot, which resulted in 72 knots (or 1 knot every 8.3 s). Scatter and background event rejection was achieved by establishing a distance threshold in the objective function of 7 mm. Cell trajectory data were plotted using MATLAB.

### *In Vivo* Cell Tracking with PET/CT

All *in vivo* procedures were approved by the Administrative Panel on Laboratory Animal Care at Stanford University under protocol 28882. Female BALB/c *nu*/*nu* mice (6 weeks old; average weight of 20 g) were used for *in vivo* studies. MDA-MB-231 cells were labeled with ^68^Ga-MSN, then different numbers of cells were injected into the animals through various routes, including intravenous, subcutaneous, intraperitoneal, and footpad injection. Radiolabeled single cells were screened using a portable radiation survey meter (Model 3 with GM detector, Ludlum, Sweetwater, TX, USA), then loaded into a syringe. After injection, static PET images were acquired (Inveon microPET/CT, Siemens pre-clinical solutions, Knoxville, TN, US; 425-650 keV energy window, 10 min acquisition). X-ray CT images were acquired using standard settings. Static PET images were reconstructed using conventional 2D-OSEM.

For dynamic tracking of single cells, a butterfly catheter was inserted into the mouse tail vein, and single radiolabeled cells were injected while PET data were acquired. The list-mode data file was imported into MATLAB and analyzed to identify the time at which the cell entered the scanner. The spline knots were then defined to achieve a higher density of knots during the initial migration of the cell and a lower density of knots after its arrest. Spline knots were defined at *T* = 85, 86.3, 87.7 and 90 s initially, then every 23 seconds thereafter. This protocol allowed us to track the cell during its rapid migration while still achieving high tracking accuracy for the remainder of the scan. Given the high density of knots in the initial phase of the trajectory, the position of the cell is approximate and should be interpreted as a qualitative estimate (see Discussion).

### *Ex Vivo* Validation Studies

To identify the organs in which single cells have arrested, *ex vivo* tissues were imaged using PET (same parameters as for *in vivo* imaging) and radioactivity was measured using gamma counting. Whole lungs were cut into small chunks and individually measured with the gamma counter. Finally, the tissue chunks were imaged using a SteREO Discovery V20 microscope (Carl Zeiss, Thornwood, CA, USA) and Zeiss LSM-700 confocal microscope (Carl Zeiss, Thornwood, CA, USA) to confirm fluorescence of DiO/DiI-labeled cells in *ex vivo* tissues.

### Statistical Analysis

All data are presented as mean ± standard deviation. P values < 0.05 were considered to be statistically significant and were obtained using Student’s *t*-test (two-tailed, unpaired samples).

### Data availability

Raw PET/CT data (list-mode format) that support the findings of this study are available from the corresponding author upon request. All other data are available within the paper and its supplementary information files.

### Code availability

Computer codes used to reconstruct cell trajectories from list-mode PET data are provided as supplementary data.

