## Supplementary information for "CellGPS: Whole-body tracking of single cells by positron emission tomography"

### Supplementary Figures

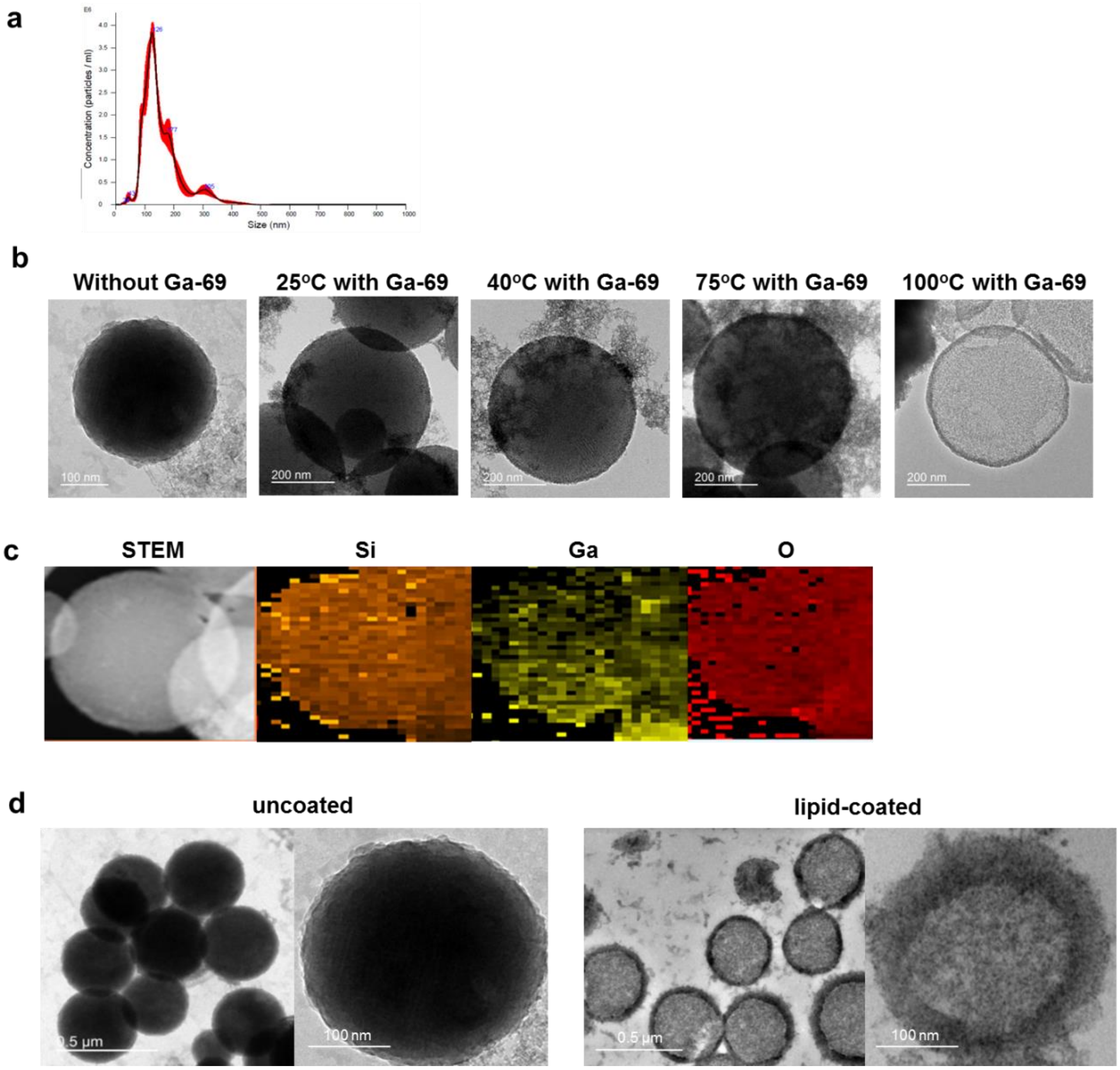


**Supplementary Figure 1. Physical characterization of MSNs.** (a) Hydrodynamic size distribution measured using Nanoparticle Track Analysis. (b) TEM images of MSNs after reaction with $^{69}$Ga for different temperatures. (c) Element mapping showing the distribution of silicon, gallium and oxygen for a small cluster of nanoparticles. (d) TEM images of MSNs before and after liposome fusion.

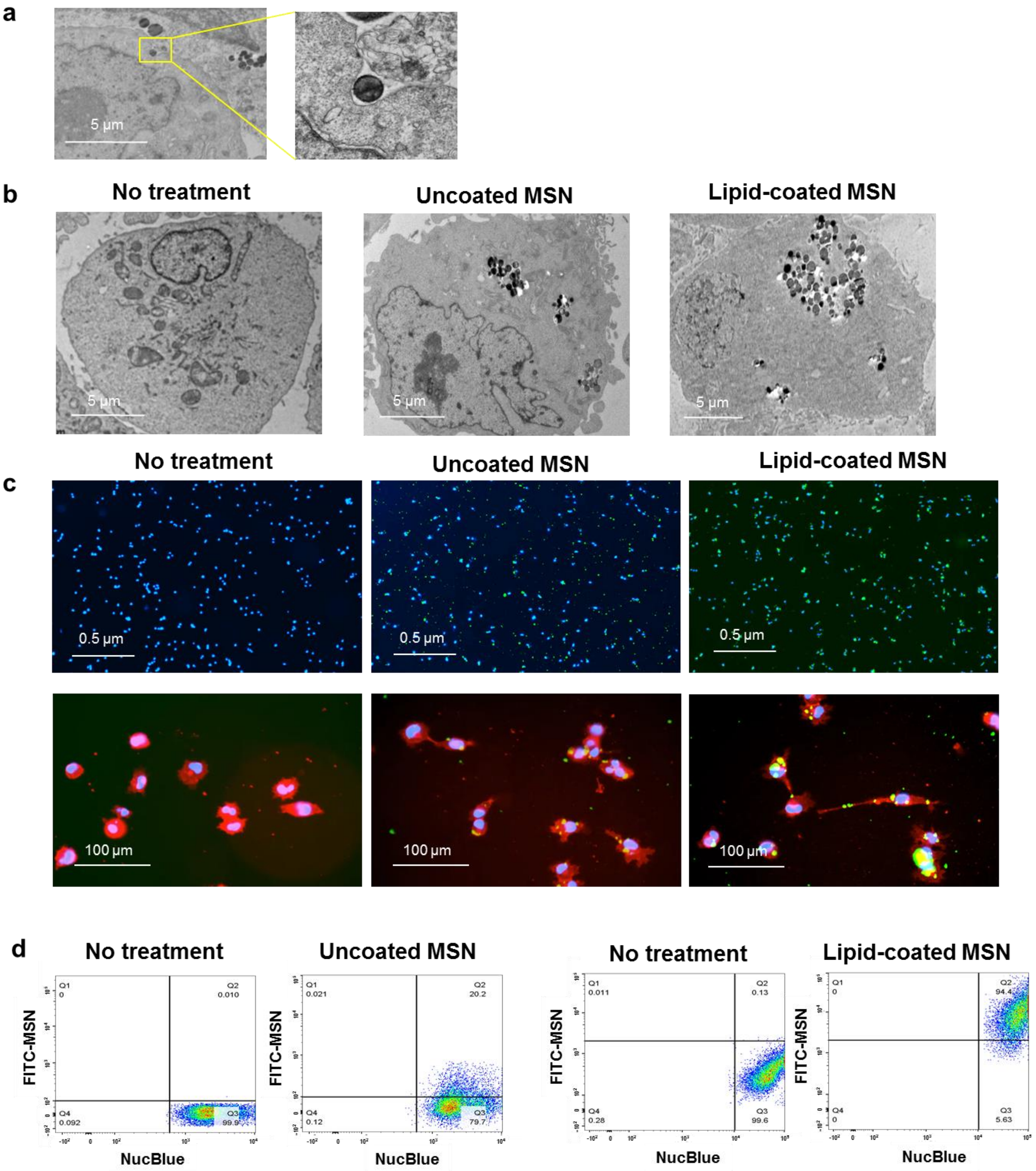


**Supplementary Figure 2. Cellular uptake of MSNs.** (a) TEM image of MSN uptake by an MDA-MB-231 cell. (b) TEM images of control cells and cells treated with uncoated and lipid-coated MSNs. (c) Fluorescence images showing cell uptake of FITC-labeled MSNs (shown in green). Nucleus is shown in blue and membrane in red. (d) Flow cytometry quantification of cellular uptake of uncoated and lipid-coated MSNs.

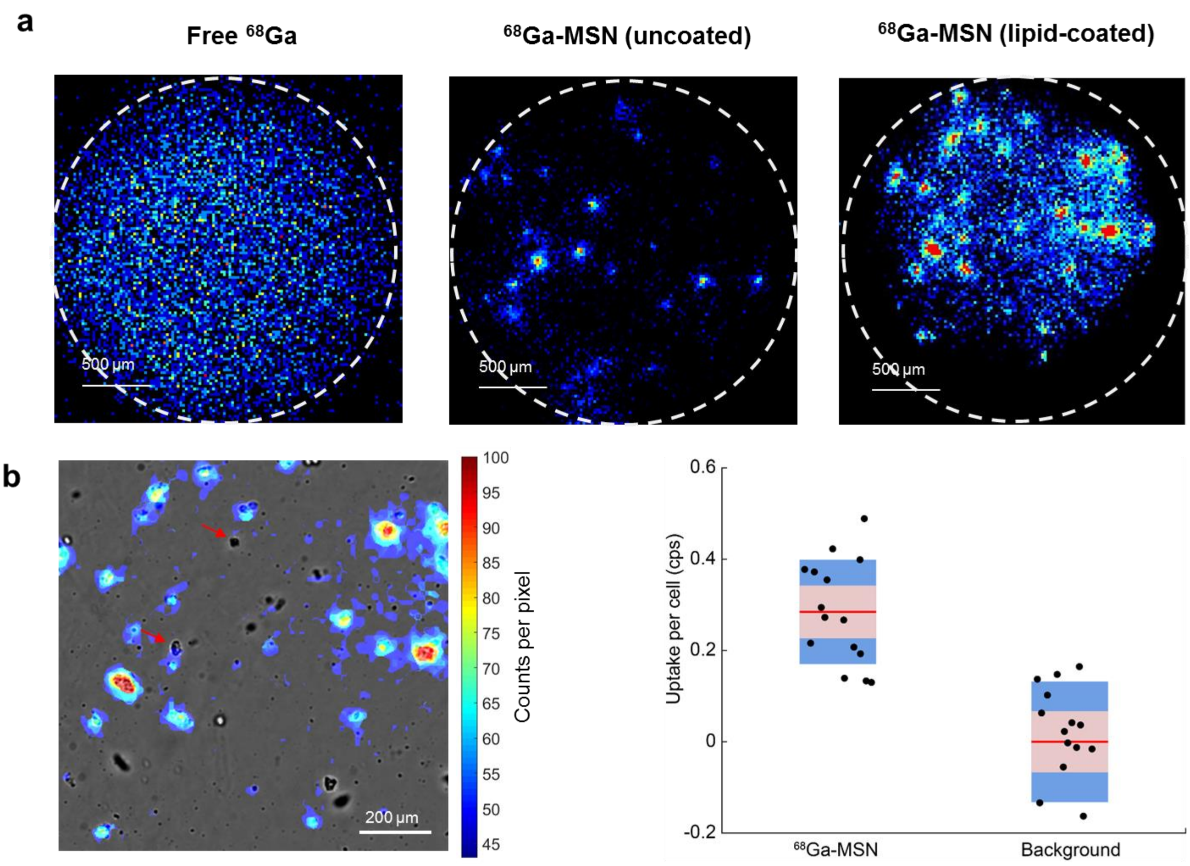


**Supplementary Figure 3. Radioluminescence microscopy (RLM) imaging of $^{68}$Ga-MSN uptake by live MDA-MB-231 cells.** (a) RLM images for cells treated with free $^{68}$Ga, $^{68}$Ga-MSN and lipid-coated $^{68}$Ga-MSN. The dashed circle outlines the field of view of the microscope. Higher background is observed when using lipofectamine for coating the MSNs. (b) RLM image showing uptake of lipid-coated $^{68}$Ga-MSN, overlaid on brightfield image for comparison. Region-of-interest analysis was used to quantify $^{68}$Ga uptake per cell in counts per second (cps). A few cell clusters (up to four cells/cluster) were included in the analysis by dividing the uptake by the number of cells. High background was measured, possibly due to non-specific binding of the lipid-coated MSNs to the glass-bottom imaging dish. Cell-to-cell variability was consistent with background variability, suggesting relatively homogenous uptake of the MSNs by cells. A few cells (red arrows) had no detectable uptake.

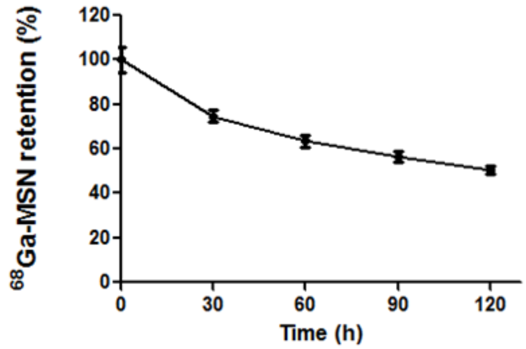


**Supplementary Figure 4.** $^{68}$Ga-MSN efflux from MDA-MB-231 cells, as a function of time. Cells were labeled with $^{68}$Ga-MSN according to the standard protocol. The cells were washed at different timepoints and retrieved for gamma counting to assess $^{68}$Ga retention. Experiments performed in triplicate.

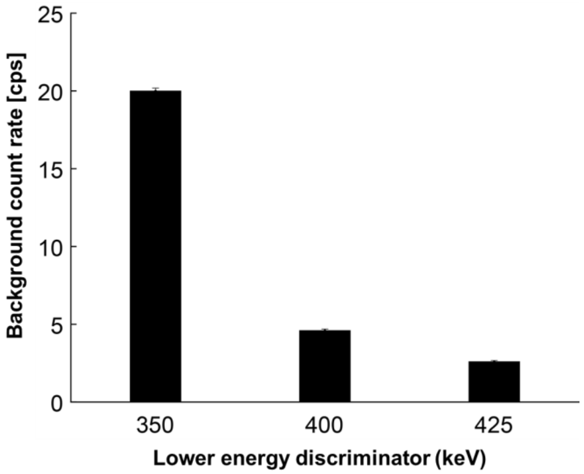


**Supplementary Figure 5.** Background count rate of Inveon PET/CT scanner for different energy window, measured with no activity in the field of view. Upper energy discriminator was 650 keV. Scan time was 10 min with no activity in the field of view.

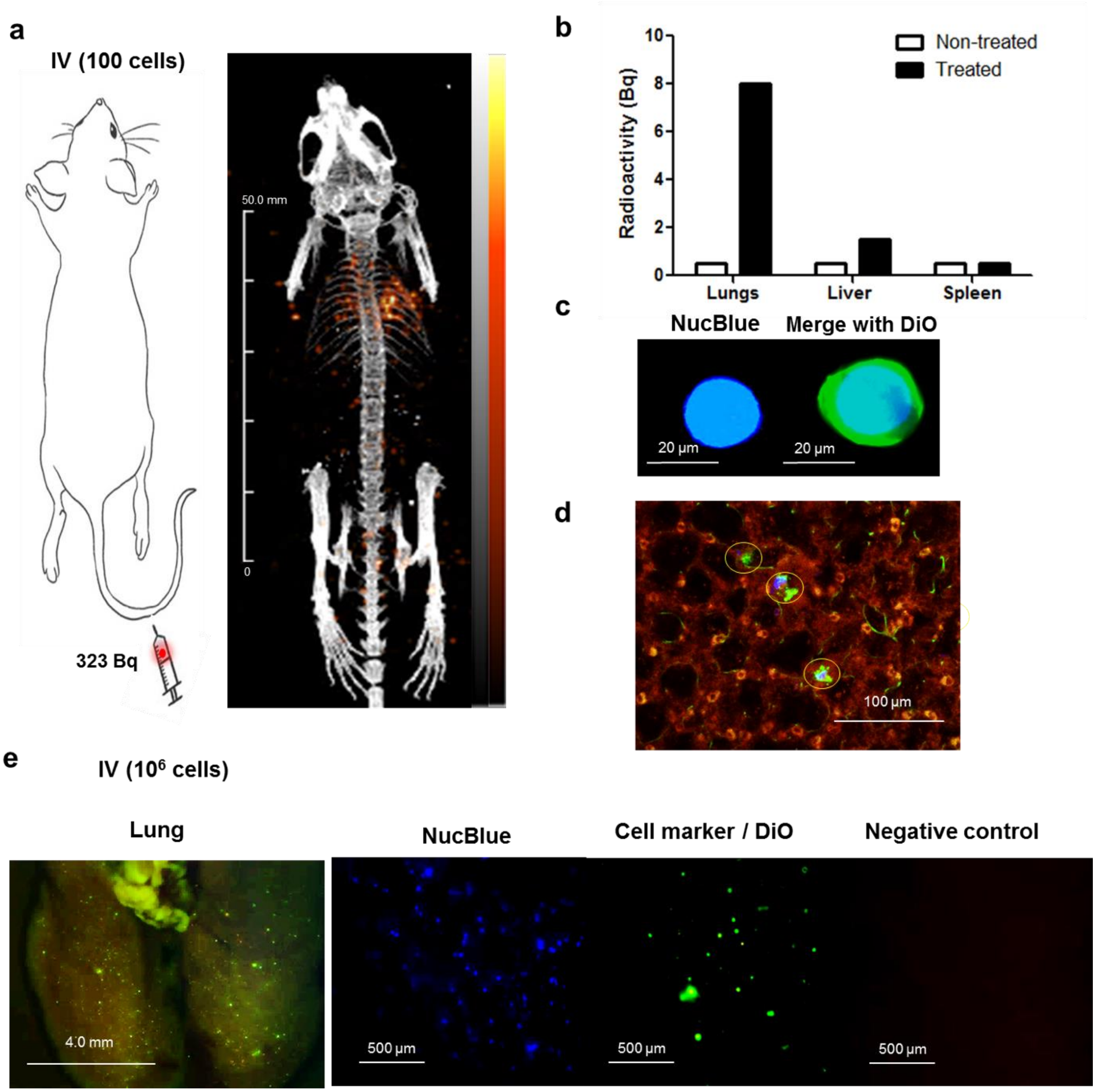


**Supplementary Figure 6. *In vivo* PET imaging of IV-injected cells.** (a) Whole-body PET/CT acquired minutes following IV injection of 100 radiolabeled cells. (b) *Ex vivo* organ radioactivity quantified by gamma counting. (c) Fluorescence imaging of single MDA-MB-231 cell following staining with NucBlue and DiO. (d) *Ex vivo* fluorescence imaging shows arrest of injected cells in the lungs (blue: NucBlue, green: DiO, red: autofluorescence). (e) *Ex vivo* fluorescence imaging shows arrest of $10^6$ injected cells in the lungs. Negative control is shown by imaging red fluorescence.

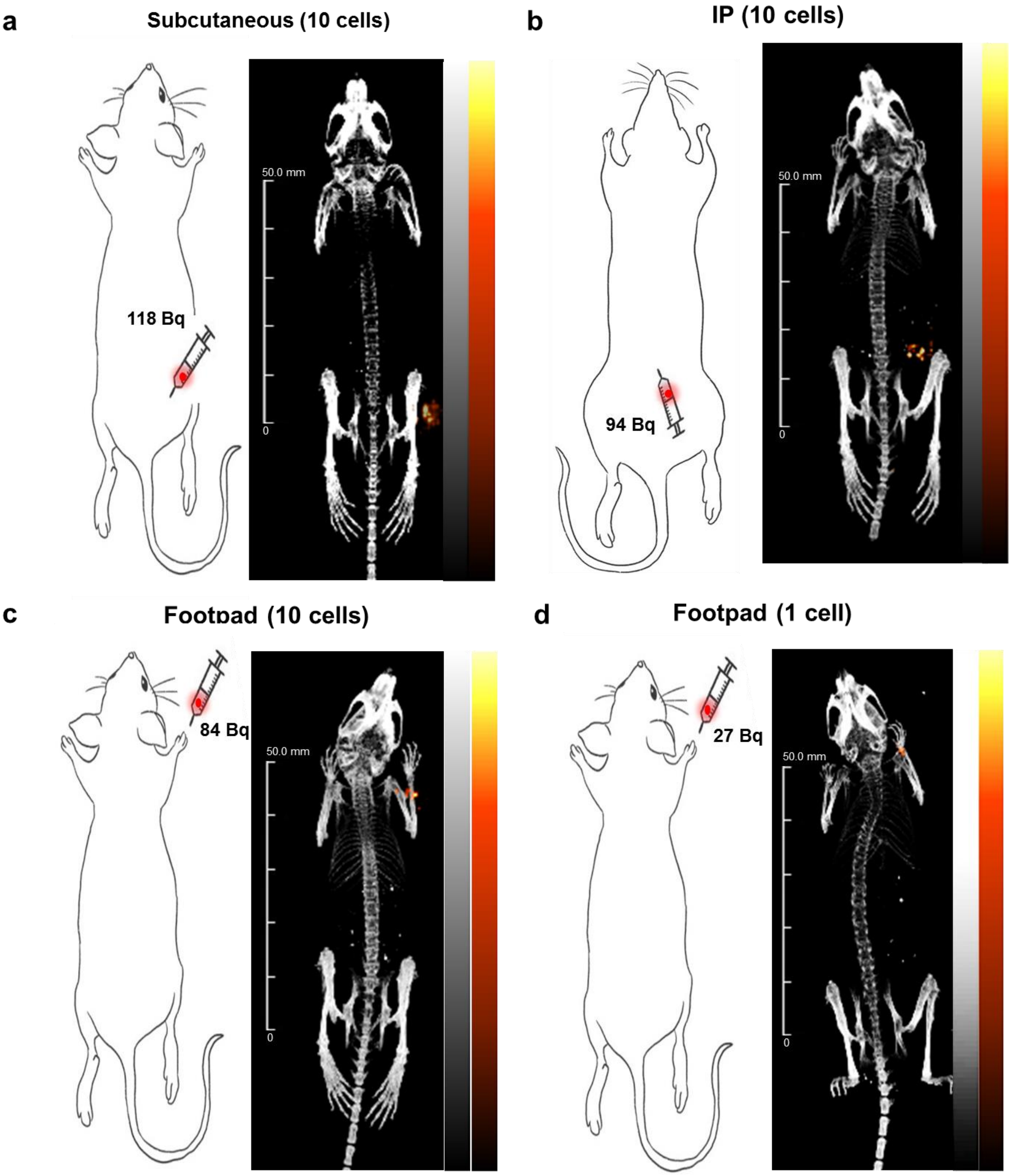


**Supplementary Figure 7. Static whole-body PET imaging of small number of injected cells in mice.** (a) Whole-body PET image after subcutaneous injection of radiolabeled 10 cells. (b) Same, after intraperitoneal injection. (c) Same, after injection in the footpad. (d) Whole-body PET imaging of a single radiolabeled cell after injection in the footpad.

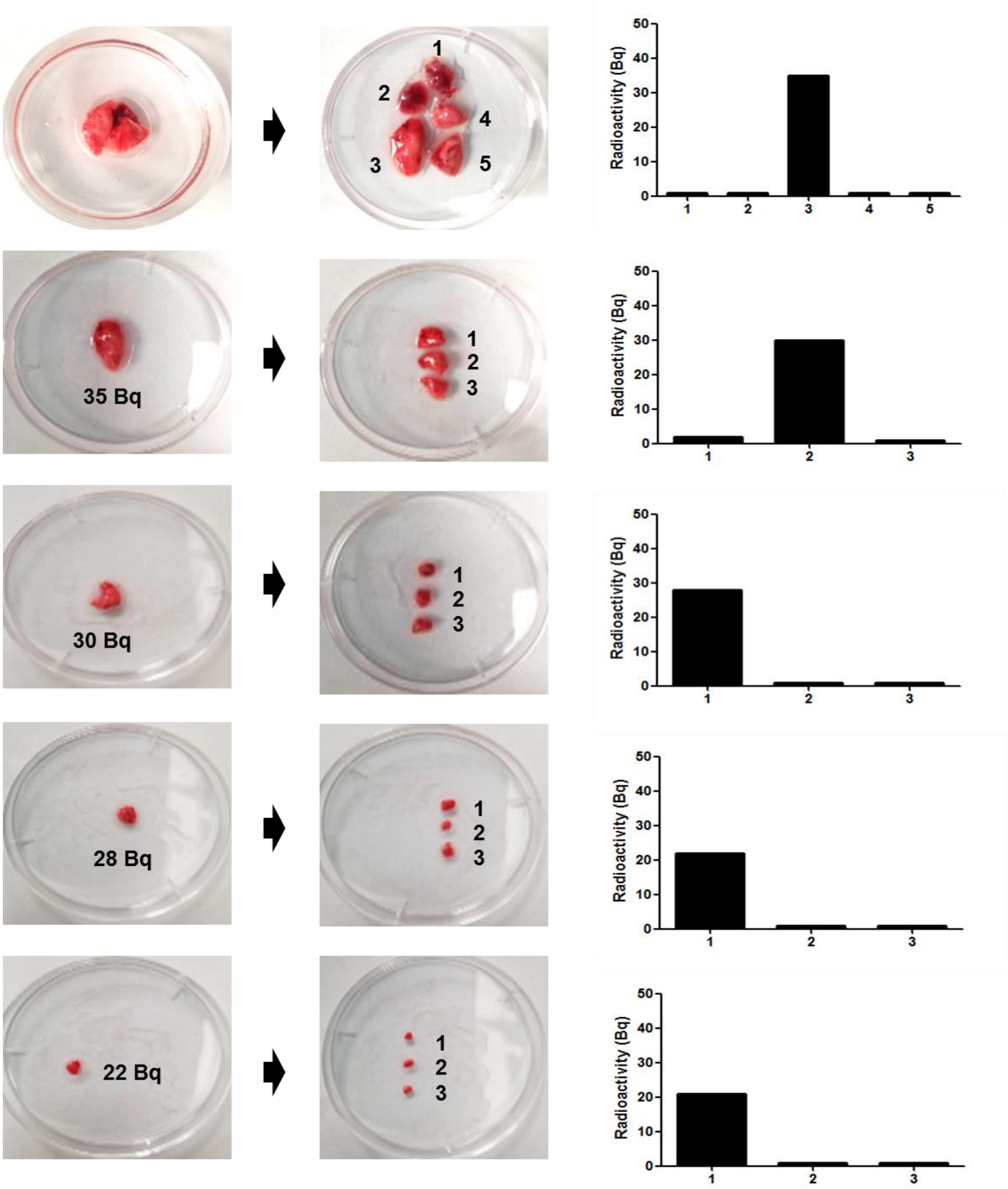


**Supplementary Figure 8.** Isolation of a single radioactive cell from lung tissue. Radioactive tissue specimens were repeatedly cut into smaller chunks until the radioactive tissue was no larger than 1 mm.

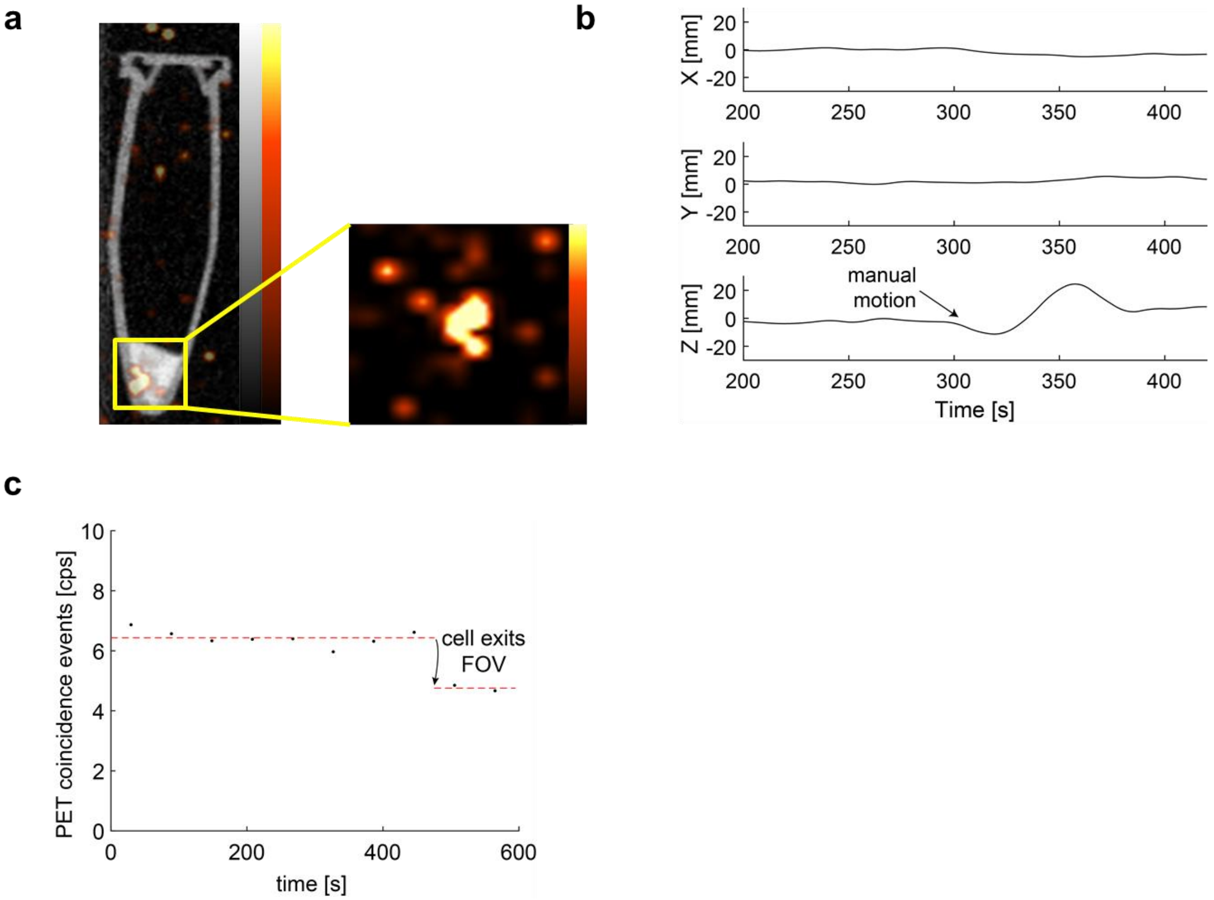


**Supplementary Figure 9. Dynamic tracking of a radiolabeled single cell using PET/CT.** (a) A single radiolabeled cell (36 Bq) was placed in a 1.5 ml conical tube and imaged by PET/CT. (b) Cell trajectory reconstructed the vial was moved by an operator inside the bore of a PET scanner (Inveon D-PET, energy window 400-650 keV). (c) Coincidence event rate recorded by the PET scanner before and after the vial was removed from the field of view. The background coincidence rate is due to intrinsic radioactivity of the PET detectors.

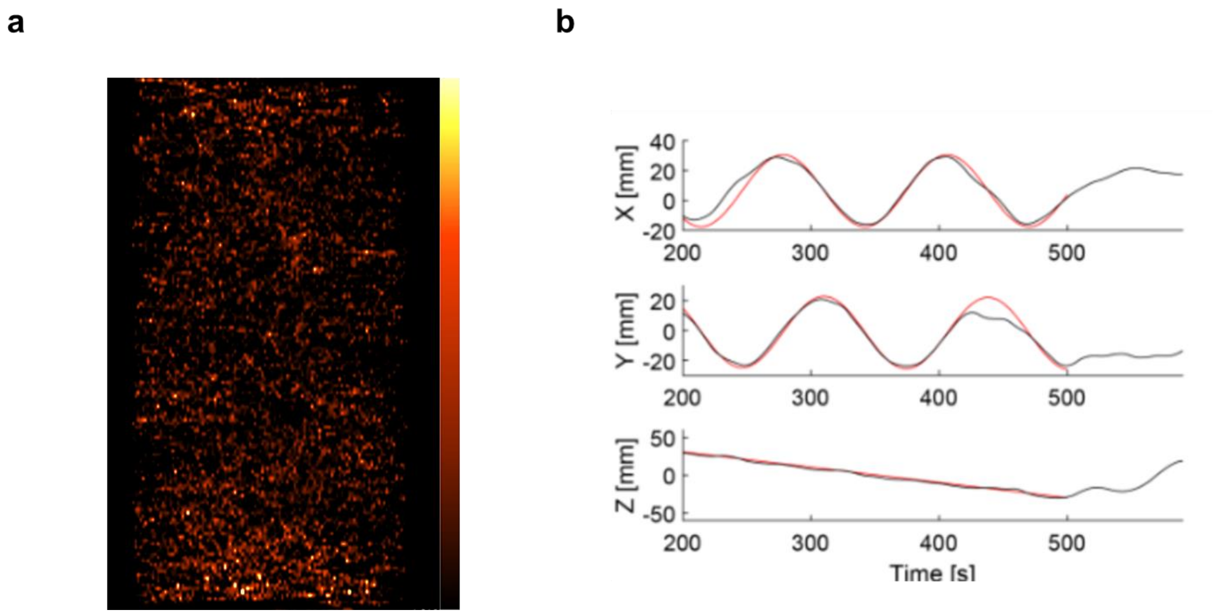


**Supplementary Figure 10. Dynamic PET tracking of single cell flowing through helical phantom.** (a) Static PET image, reconstructed using conventional OSEM, of a single radiolabeled cell (67 Bq) moving through a length of plastic tubing coiled around a 3D printed cylinder. The trajectory of the cell cannot be seen in this image. (b) Same data, but reconstructed using spline-based reconstruction algorithm, clearly highlights the positon of the single cell as a function of time.

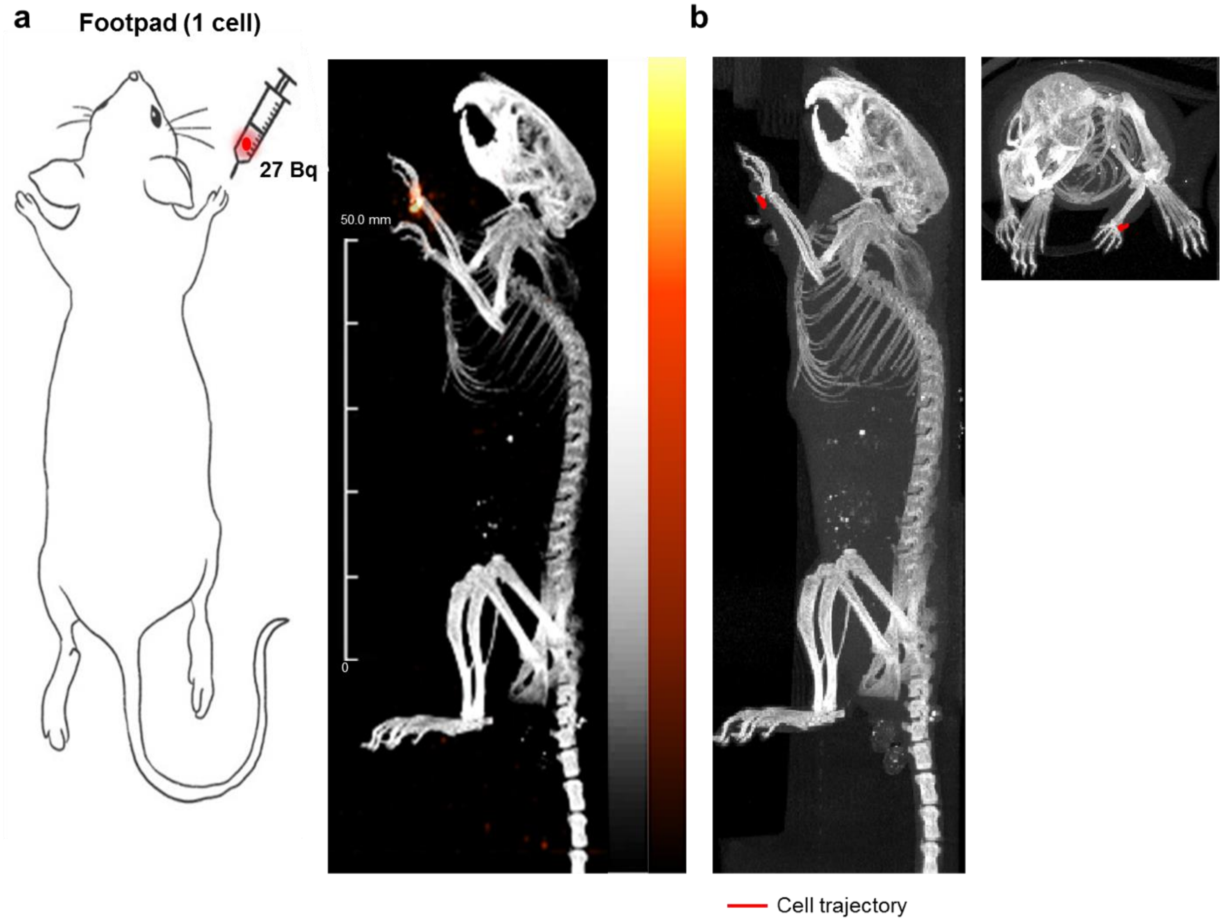


**Supplementary Figure 11. Dynamic PET tracking of single cell following footpad injection.** (a) Conventional PET/CT reconstruction shows single cell in injection site after injection in the fore paw (previously shown as Fig. 3c). (b) Dynamic trajectory reconstructed from the same list-mode data localizes the single cell to the injection site. Two orthogonal views are shown to confirm localizations. (1,362 events, 14 spline knots)

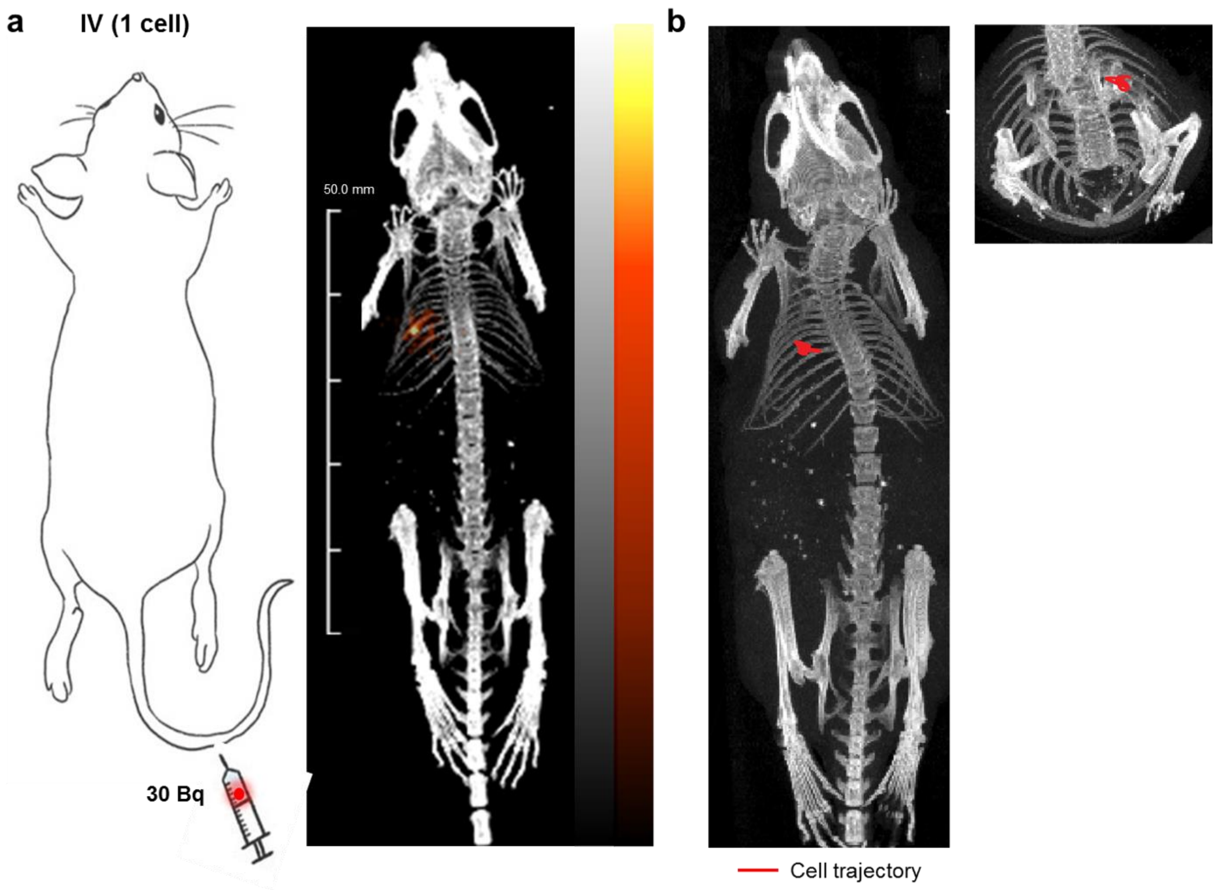


**Supplementary Figure 12. Dynamic PET tracking of single cell following IV injection.** (a) Conventional PET/CT reconstruction shows single cell in lung after IV injection (previously shown as Fig. 3d). (b) Dynamic trajectory reconstructed from the same list-mode data localizes the single cell to the lung. (1,643 list-mode events, 16 spline knots)

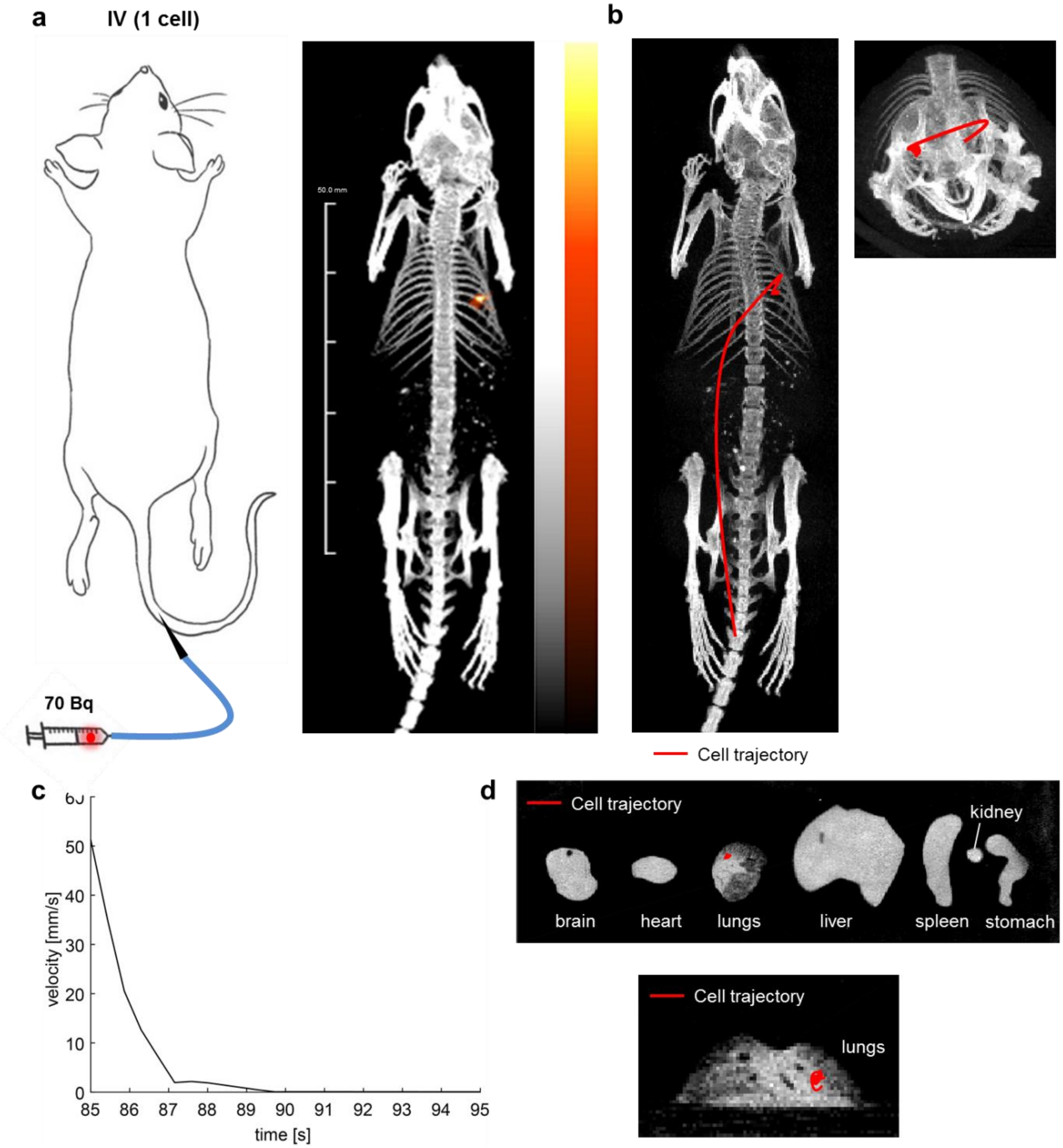


**Supplementary Figure 13. Dynamic PET tracking of a single cell, during and after IV injection.** (a) A single cell (70 Bq) was injected via catheter into the tail vein. Conventional OSEM reconstruction localized the single cell in lung, where it had arrested. The PET acquisition started approximately 1 min before the cell was injected. (b) Dynamic trajectory reconstructed from the same list-mode data tracks the single cell as it travels through the bloodstream and arrests in the lungs (10,790 events in total, 25 spline knots, 10 min). (c) Velocity of the injected cells, estimated from the reconstructed trajectory. (d) Dynamic cell trajectory, reconstructed from a PET/CT acquisition of the excised organs (7,188 events, 24 spline knots, 10 min). The fluctuation in the position of the single cell is due to noise and statistical uncertainty.

### Supplementary Video Captions

**Supplementary Video 1.** Cell position (red dot) reconstructed from list-mode PET data, over time. A single cell (36 Bq) was placed in a 1.5 ml conical tube and imaged by PET/CT. The vial containing the cell is moved between time points $t$=300 and $t$=380 seconds. The blue lines represent the coincidence events recorded by the scanner. The black line is the cell trajectory.

**Supplementary Video 2.** Cell position (red dot) reconstructed from list-mode PET data, over time. A single cell (67 Bq) was flown moving through a length of plastic tubing coiled around a 3D printed cylinder. The blue lines represent the coincidence events recorded by the scanner. The black line is the cell trajectory.

**Supplementary Video 3.** Cell position (red dot) reconstructed from list-mode PET data, over time. A single cell (27 Bq) was injected into the forepaw of a mouse. The blue lines represent the coincidence events recorded by the scanner. The yellow line is the cell trajectory. A CT image of the mouse in shown in the background.

**Supplementary Video 4.** Cell position (red dot) reconstructed from list-mode PET data, over time. A single cell (30 Bq) was injected into the tail-vein of a mouse. PET data were acquired a few minutes after cell injection. The blue lines represent the coincidence events recorded by the scanner. The yellow line is the cell trajectory. A CT image of the mouse in shown in the background.

**Supplementary Video 5 and 6.** Cell position (red dot) reconstructed from list-mode PET data, over time. A single cell (70 Bq) was injected into the tail-vein of a mouse via a catheter, while PET data were acquired. The blue lines represent the coincidence events recorded by the scanner. The yellow line is the cell trajectory. A CT image of the mouse in shown in the background.

**Supplementary Video 7.** Cell position (red dot) reconstructed from list-mode PET data, over time. A single cell (70 Bq) was injected into the tail-vein of a mouse via a catheter, then the organs were excised and imaged with PET/CT. The blue lines represent the coincidence events recorded by the scanner. The yellow line is the cell trajectory. A CT image of the *ex vivo* in shown in the background.
